# *Wnt* and *Nodal* asymmetries stratify mouse laterality phenotypes in the absence of node flow

**DOI:** 10.64898/2025.12.21.695689

**Authors:** Amaia Ochandorena-Saa, Emeline Perthame, Zoé Oulerich, Alexander Chamolly, Thierry Blisnick, Johanna Lokmer, Cécile Rouillon, Philippe Bastin, Sigolène M. Meilhac

## Abstract

Left-right symmetry breaking in mice is considered to occur via node leftward fluid flow. The molecular cascade within the node has been reconstructed at high resolution. However, its importance for organogenesis remains poorly understood. Here we show in mutants for the motile cilium component CCDC40 that 70% of mice develop normal situs at birth despite abrogation of node flow. The discrete morphospace output, including situs inversus totalis and heterotaxy with left isomerism, supports a novel model of symmetry-breaking, in which node flow only biases asymmetry orientation, while another mechanism, potentially self-amplifying, generates it. Longitudinal, quantitative and paired transcriptomic analyses uncover the molecular signature of laterality clusters, highlighting WNT in addition to NODAL pathway asymmetry. We identify asymmetry of cardiopulmonary progenitors, the disruption of which is associated with combined heart and lung defects in mutants. Our work refines the role of the node in the establishment of asymmetry.

## Introduction

Left-right patterning of visceral organs is not just an anatomical curiosity but a functional requirement. Defective coordination of left-right patterning leads to the severe syndrome of heterotaxy (or situs ambiguus), characterised by abnormal organ shapes or discordant asymmetries between organs or organ compartments. Heterotaxy affects 1/10,000 live births and is frequently associated with complex congenital heart defects, which determine the prognosis of patients (*1,2*). Heterotaxy can co-segregate in families with completely normal (situs solitus) or completely mirror-image (situs inversus totalis) organ laterality, which is usually asymptomatic (*1*). Heterotaxy can be categorised as right or left isomerism, if organ compartments that ought to be asymmetric are instead similar (isomeric) on both sides, for example lung lobes, spleen and atria. However, heterotaxy encompasses a broad spectrum of defects, with inter-individual variations.

Despite phenotypic variations, a monogenetic origin has been identified in about 20% of heterotaxy cases (*3*). Patient genome sequencing efforts have currently uncovered 42 distinct OMIM genes associated with heterotaxy. Many were identified based on knowledge acquired in the mouse model of the mechanisms establishing asymmetry in the early embryo. The TGFb secreted factor NODAL was first discovered as a major left determinant (*4–6*), and its inactivation impairs the morphogenesis of visceral organs, including the heart, lungs, spleen, stomach, liver and intestine, thus modelling heterotaxy with right isomerism (*7–9*). The laterality of *Nodal* expression was found abnormal in a spontaneous mouse mutant (*Dnah11^iv^*) and a mutant arising from a transgenic insertion (*Invs^inv^*), showing that the underlying ciliary genes are upstream determinants of laterality (*6*). A breakthrough came with the discovery of motile cilia in a pit of cells referred to as the node, establishing a leftward fluid flow as the earliest event of symmetry breaking in mammals (*10,11*). The node is now considered as a left-right organiser (*12*). Forcing node flow in a rightward direction by microfluidics can reverse the expression laterality of the NODAL target *Pitx2* (*13*). The flow may carry chemical cues (*14,15*) or can be sensed mechanically by primary cilia of node crown cells (*16,17*). Sensing of flow on the left triggers the left-sided degradation of *Dand5* mRNA, that encodes a NODAL antagonist (*18,19*). These observations have thus reconstructed a linear cascade establishing left-right patterning, from motile cilia in the node pit to node flow to elevated NODAL signaling in the left node crown, and finally activation of NODAL signaling in organ precursor cells in the left lateral plate mesoderm (*20–22*).

Cilia are organelles built on a scaffold of microtubule doublets or axoneme. Motility is provided by DYNEIN motor proteins arranged as inner and outer arms. These arms are assembled along the axoneme with a 96nm periodicity, set by a molecular ruler complex, composed of CCDC39 and CCDC40 (*23,24*). Thus, *Ccdc40* is required for ciliary beat. In humans, it is the third most frequent gene associated with the respiratory disease primary ciliary dyskinesia, and is also associated with infertility and situs inversus totalis (*25–27*). In the mouse, the *lnks* allele of *Ccdc40* has an ENU-induced nonsense point mutation, which introduces an early stop codon, and models motile ciliopathy, primary ciliary dyskinesia and laterality defects (*25*). Within the framework of a linear cascade of left-right patterning, paralysed cilia are predicted to abrogate flow and maintain bilateral *Dand5* expression. However, in mutants for motile cilium genes *Dnah11*, *Dnah5* and *Ccdc40*, all possible patterns of laterality have been observed, including absent, bilateral or unilateral (left or right) NODAL signaling, overall reported as randomisation of left-right asymmetry (*6,28,29*). The origin of this variability is poorly understood. Does it reflect truly random mechanisms of symmetry breaking, which are normally overridden by the node flow? Could variability arise from experimental pitfalls, which do not provide a complete picture of the mutant phenotype? It has been proposed that two cilia are sufficient to generate enough flow and trigger the asymmetry cascade (*30*), suggesting that variability in flow intensity could underlie variable laterality outputs in mutants. Alternatively, asymmetric gene expression is very transient (*9,31*), so that absent expression could reflect an inappropriate stage of observation. In keeping with this, NODAL signaling has been reported to be delayed in motile cilium mutants (*28,29,32*). Until now, it has not been possible to assess the phenotypic output of a given pattern of *Nodal* expression in the early embryo, because it requires observations at different stages and partial penetrance precludes correlation. When laterality was diagnosed at fetal stages, based on organs, 18-20% of mutants for *Dnah5* or *Dnai1* displayed normal situs solitus, whereas the rest have situs inversus totalis or heterotaxy. However, the developmental trajectory of these fetuses is unknown. Thus, the role of the left-right organiser has remained unclear.

Asymmetric cues other than *Nodal* have been identified such as *Wnt3* in the left node crown (*33,34*), and BMP signalling in the right lateral plate mesoderm (*35*). However, within the framework of a linear cascade of left-right patterning, only NODAL signaling is routinely assessed in laterality mutants, leaving the question open of whether these alternative asymmetric factors are mainly required in feedback loops to reinforce NODAL signaling or for independent functions.

The heart provides an interesting readout of laterality mechanisms. It is asymmetric in its function, driving a double blood circulation. Heart morphogenesis is sensitive to left-right patterning (*36*). This is first detected by the rightward looping of the embryonic heart tube. However, heart looping is more than just a direction, usually scored in laterality mutants; it corresponds to the transformation of an initial straight tube into a helical shape (*37*). During heart looping, the right ventricle, which was lying cranially, is repositioned on the right side of the left ventricle. We have previously shown that heart looping is centered on a buckling mechanism, when the heart tube grows between fixed poles (*38*). Tube elongation is fed by different populations of precursor cells located dorsally, ventrally or posteriorly (*39*). *Nodal* is transiently expressed in left cardiac precursors and controls independent asymmetries at the arterial and venous poles. *Nodal* is not required to initiate asymmetric morphogenesis, but to bias and amplify buckling (*9,40*). Our analysis of heart morphogenesis thus shows that left-right patterning is not a simple linear cascade, but rather entails several processes contributing to asymmetry. It is a current challenge to unravel the contribution of individual asymmetric cues to organogenesis. We have developed a framework for a comprehensive analysis of laterality defects, including 3D imaging and quantification, at anatomical and molecular levels (*9,40,41*). We have also established a paired transcriptomic approach, able to uncover asymmetric gene expression in the lateral plate mesoderm and avoiding noise from inter-individual and stage variations (*40*). This has opened the possibility to investigate with higher precision how cells perceive asymmetry during organogenesis.

Here we analyse the role of the node, using *Ccdc40^lnks^*mutants as a model of motile ciliopathy. We perform in-depth analysis of laterality defects, using quantitative 3D imaging, showing that the left-right organiser is dispensable in 70% of cases to establish organ laterality. We tackle the challenge of correlating molecular asymmetries with measure of the node flow on one hand, and with morphological phenotype on the other hand. Unsupervised clustering of samples identifies key geometrical parameters and domains of NODAL signaling as signatures of heterotaxy compared to situs solitus or situs inversus totalis. Finally, we provide broader profiling of molecular asymmetries based on paired transcriptomics. This reveals the WNT pathway as an important determinant to stratify laterality phenotypes. Overall, we demonstrate that the node flow is required as a bias of symmetry breaking, controlling the robustness of left-right asymmetry, but that laterality can be established without it. Our work provides novel insight into the origin of the spectrum of laterality defects.

## Results

### *Ccdc40* mutants are a model of motile ciliopathy with abrogated node flow

To understand how the motile cilium component CCDC40 affects left-right patterning, we first investigated its expression pattern. We detected high *Ccdc40* expression at E8.5b/c in the pit of the node, where motile cilia are localised (Fig. 1A-D), but not in the notochord (Fig. S1D). Co-labelling shows that *Ccdc40* is not expressed in heart progenitors in the lateral plate mesoderm (Fig. S1A-C). Published single cell transcriptomics further support specific expression of *Ccdc40* in the node (Fig. S1E). We also wondered which cell types express a toolkit of motile cilia, and interrogated single cell datasets. We found that the node/notochord was the only tissue between E6.5 and E8.5 to express a toolkit of 82 motile cilium genes (Fig. S2). We then analysed how the node is affected in *Ccdc40^lnks/lnks^*mutants (Fig. 1E) (*25*). By scanning electron microscopy, we found that mutant nodes had many cilia with an abnormal ballooned morphology (Fig. 1F-G), mainly localised in the node pit (Fig. S1F-H). We evaluated their function by live imaging of fluorescent bead movements. Compared to controls, mutant nodes were unable to generate a detectable flow (Fig. 1H-J, Movie S1). We conclude that *Ccdc40* mutants are a model of motile ciliopathy with fully penetrant abrogated node flow.

**Fig. 1.**
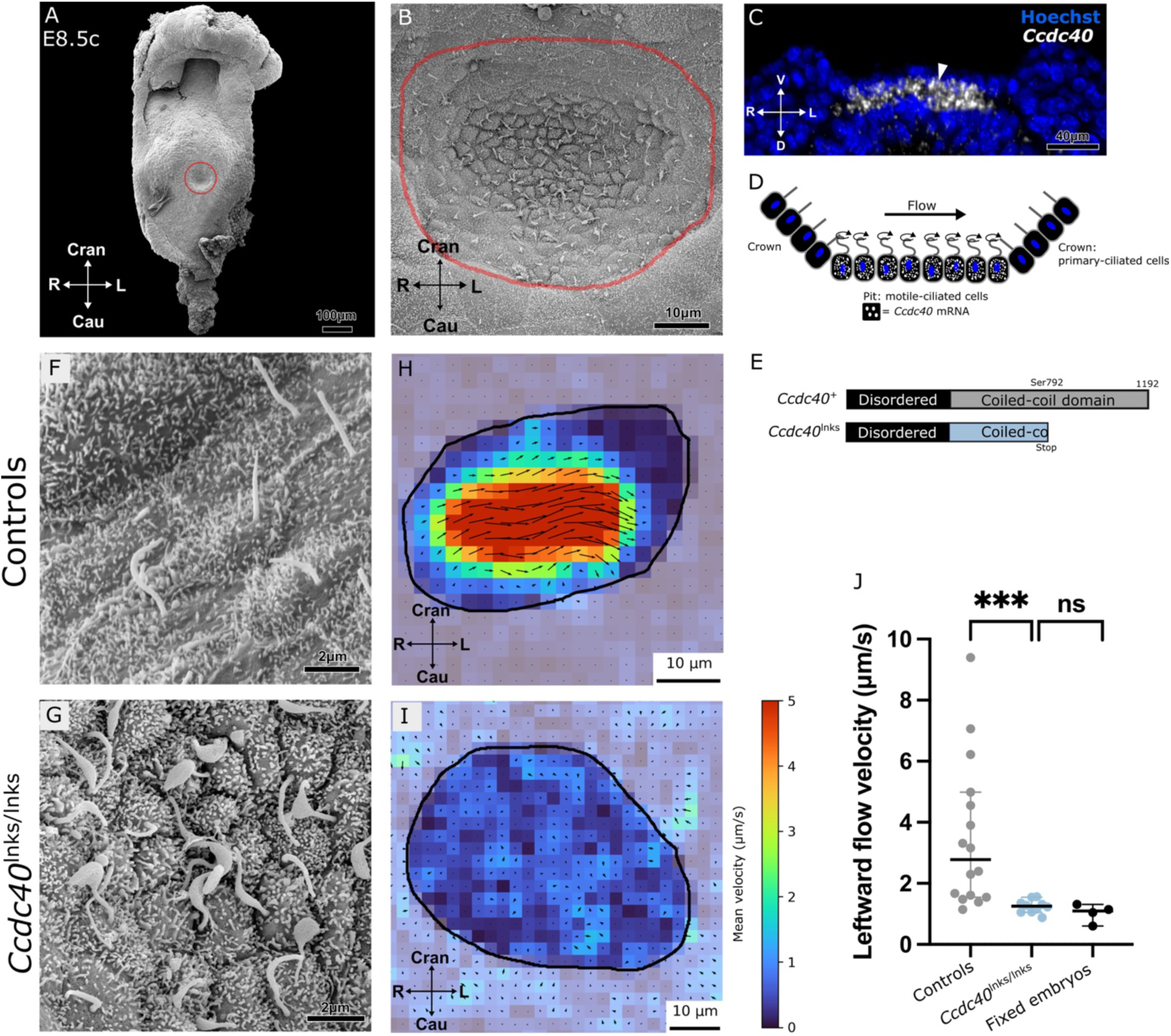
*Ccdc40* mutation disrupts the shape and function of motile cilia in the node pit. (**A-B**) Images by scanning electron microscopy of an E8.5c wild-type mouse embryo seen on a ventral view. The node is outlined in red and shown at higher magnification in (B). (**C**) Transverse optical section of the node, counterstained with Hoechst, showing *Ccdc40* expression (white) in the pit (white arrowhead) by whole mount in situ hybridisation. (**D**) Schema of the node. (**E**) Predicted CCDC40 protein domains produced in wild-type (+) and mutant (lnks) alleles. (**F-G**) High magnification of cilia in the node pit of control (F, n=3) or *Ccdc40^lnks/lnks^* mutant (G, n=3) embryos, imaged by scanning electron microscopy. (**H-I**) Map of the node flow by Particle Image Velocimetry of fluorescent bead movements in control (H) or *Ccdc40^lnks/lnks^* (I) live embryos at E8.5b. (**J**) Corresponding quantification of flow velocity at E8.5b-d. Means and standard deviations are shown. *** p value < 0.001; ns, non-significant (Mann-Whitney test, n=16 *Ccdc40^+/+^* or *Ccdc40^lnks/+^* control live embryos, n=10 *Ccdc40^lnks/lnks^* mutant live embryos, n=4 control fixed embryos). Cran, cranial; Cau, caudal; D, dorsal; R, right; L, left; V, ventral. See also Fig. S1-S2, Table S6 and Movie S1.

### The node is required for the robustness of visceral organ laterality

We analysed laterality defects upon loss of *Ccdc40* by 3D imaging of visceral organs in situ at birth. Based on the clinical nomenclature, we found three categories of laterality phenotypes, with no sign of in utero lethality (Fig. 2A). In contrast to situs solitus in controls (Fig. 2B, Movie S2), 15% of *Ccdc40* mutants had situs inversus totalis, in which visceral organs are in mirror-image, as well as the heart structure (Fig. 2D, Movie S2). We did not detect congenital heart defects associated with situs inversus totalis, but some mild anomalies in lung or liver lobation, or in the heart apex position (Table S1). Another 15% of mutants had heterotaxy with left isomerism of the lungs and bronchi (Fig. 2E-F, Movie S2). The heart always had complex defects, in the venous return, atrial appendages (100% left isomerism), ventricle laterality (50% D-loop, 50% L-loop), septation, ventriculo-arterial connections, position of the great arteries or aortic arch (Table S1). Two cases also had laterality defects in the stomach, spleen, liver and colon. Surprisingly, the majority of mutants (70%) had no laterality defects, with either complete situs solitus (Fig. 2C, Movie S2) or mild asymptomatic anomalies in lung or liver lobation, in the trajectory of the inferior caval vein or in the colon flexure (Table S1). Thus, despite the full penetrance of node disruption, *Ccdc40* mutants display partial penetrance of laterality defects at birth. The distribution of the three groups of laterality phenotypes significantly differs from a uniform randomisation hypothesis (p=0.0002, chi-square test). This shows that the node is dispensable to initiate asymmetric organogenesis, and only required for the robustness of laterality patterning.

**Fig. 2.**
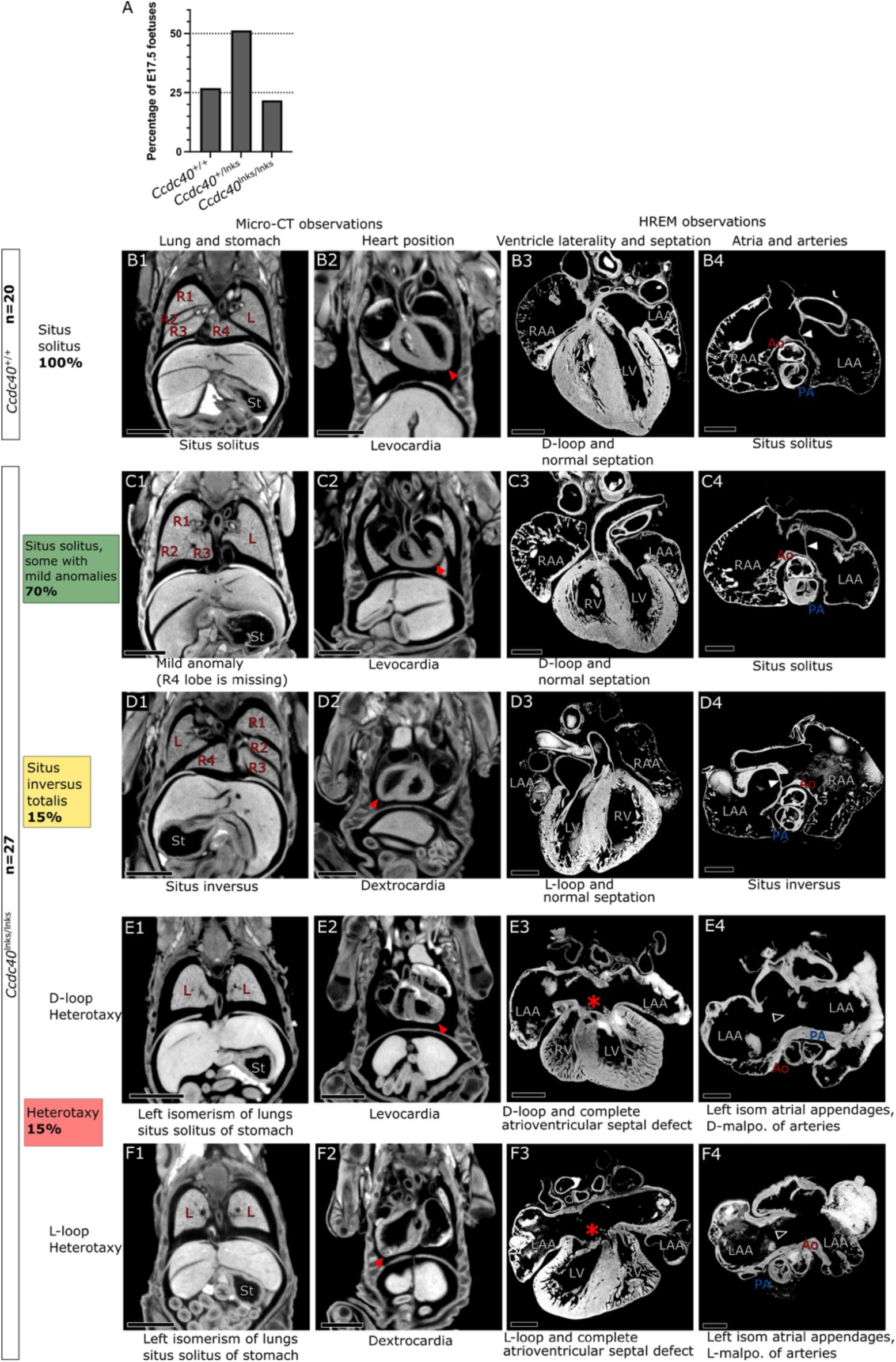
*Ccdc40* mutants display partial penetrance of laterality defects at birth. (**A**) At E17.5, *Ccdc40* mutants were found at the expected Mendelian ratio (p=0.703, chi-square test; n=115). Examples of control (**B**) and *Ccdc40* mutants (**C-F**) at E17.5 or P0. (B1-F1, B2-F2) Frontal sections from 3D images by micro-CT showing the position of visceral organs in the thoracic and abdominal cavities. Red arrowheads point to the heart apex. (B3-F3, B4-F4) Frontal (B3-F3) and transverse (B4-F4) sections of explanted hearts imaged in 3D by High Resolution Episcopic Microscopy (HREM). The asterisk indicates complete atrioventricular septal defect. White arrowheads point to the presence (filled arrowhead) or absence (empty arrowhead) of atrial septation. *Ccdc40* mutants have partial penetrance of laterality defects: situs inversus totalis (D) or heterotaxy with left isomerism of the airways (E-F). Scale bars: 2000 μm (B1-F1, B2-F2), 500 μm (B3-F3, B4-F4). Sample size (n) refers to number of foetuses in each category. R1-R4, right lung lobes; L, left lung lobe; St, stomach; LAA, left atrial appendages; RAA, right atrial appendages; LV, left ventricle; RV, right ventricle; Ao, aorta; PA, pulmonary artery. See also Tables S1, S6 and Movie S2.

### Embryonic heart looping can be defective in *Ccdc40* mutants

The heart is the first organ to undergo asymmetric morphogenesis. We investigated the phenotype of *Ccdc40* mutants at E9.5, when looping of the embryonic heart tube is complete. In controls, the embryonic heart forms a rightward helix. In most mutants, heart looping was also rightward. However, it was inverted, i.e. leftward, in 27% of cases (Fig. 3A-C). Because heart looping can be affected in its shape and not just in its direction (*9,38*), we segmented and quantified it (Fig. 3D-H, Movie S3). The length of the heart tube in mutants was not significantly changed, indicating that heart growth is normal (Fig. 3F). However, we found occasional defects in the orientation of the right ventricle-left ventricle axis, which failed to align perpendicular to the notochord (4/50 mutants, Fig. 3D, G). Mutant hearts also showed anomalies in the position of the atrioventricular canal, which was significantly more medial (7/50, Fig. 3E, H). Overall, our quantifications at E9.5 indicate three categories of heart looping in *Ccdc40* mutants: normal, complete mirror-image and defective. Defective looping was observed in both rightward and leftward heart tubes. This parallels the laterality phenotypes at birth, situs solitus, situs inversus totalis and heterotaxy with D-loop or L-loop.

**Fig. 3.**
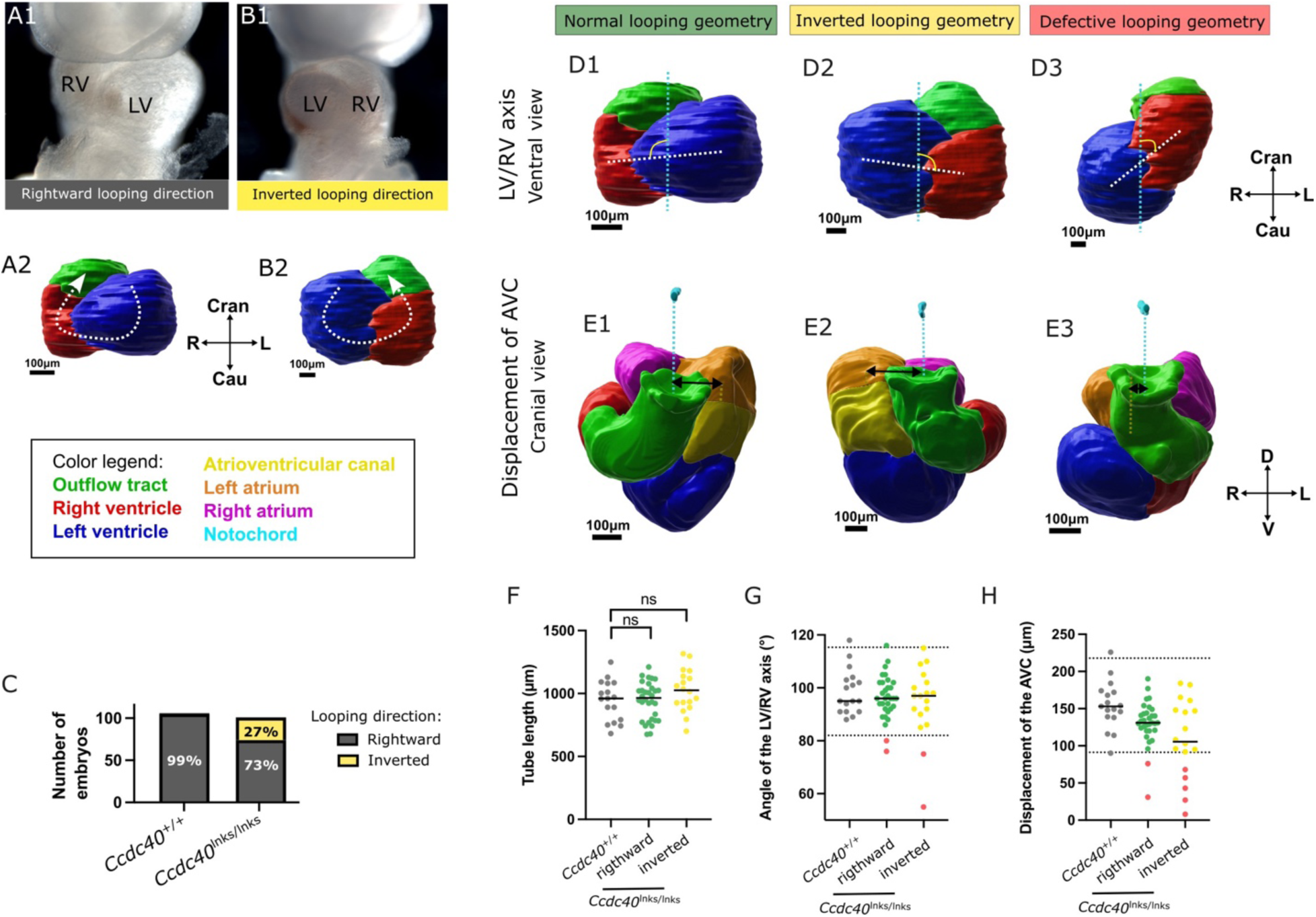
Quantification of heart looping defects in *Ccdc40* mutants at E9.5. (**A-B**) Brightfield images (A1-B1) of *Ccdc40* mutant embryos at E9.5, showing rightward or leftward (inverted) heart looping direction. (A2-B2) 3D segmentation of the embryonic heart tube with the looping direction outlined by a dashed arrow. (**C**) Bar diagram showing the frequency of heart looping direction (n=106 controls and 101 *Ccdc40* mutants). (**D-E**) 3D segmentation of the embryonic heart tube in *Ccdc40^lnks/lnks^* mutants, aligned in a ventral (D) and cranial (E) view, using the notochord as a reference axis (cyan). Examples are shown of mutants with a normal (D1-E1), mirror-image (D2-E2), or defective (D3-E3) geometry of the looped heart tube. The orientation of the RV (red) /LV (blue) axis (dashed white line) relative to the notochord (dashed blue line) is measured as an angle in (D). The lateral displacement of the atrioventricular canal (yellow) relative to the midline (dashed blue line) is measured as a distance (black double arrows) in (E). Cardiac segments are colour coded as indicated in the legend. (**F-H**) Corresponding quantifications. The tube length, which is not significantly different between controls and mutants, indicates normal growth of the heart and homogenous staging. Absolute values of the parameters in (G-H) are shown, whereas the orientation is indicated as separate rightward and inverted groups in the x-axis. ns, non-significant (t-test, n=17 *Ccdc40^+/+^* embryos, 32 *Ccdc40^lnks/lnks^*mutant embryos with rightward looping, 18 *Ccdc40^lnks/lnks^* mutant embryos with inverted looping direction). Dashed lines show the 95% distribution interval based on a Gaussian distribution of wild-type samples. 4 (G) and 7 (H) *Ccdc40^lnks/lnks^* mutants were outside this interval. Cran, cranial; Cau, caudal; D, dorsal; LV, left ventricle; R, right; RV, right ventricle; L, left; V, ventral. See also Table S6 and Video S3.

### In the absence of node flow, variable laterality of *Nodal* in the lateral plate mesoderm clusters phenotypes

Given that the node triggers the asymmetric expression of *Nodal* (*6*) and that *Nodal* regulates asymmetric heart morphogenesis (*9*), we investigated *Nodal* patterning in *Ccdc40* mutants at E8.5c. Compared to left-sided expression in the lateral plate mesoderm of controls, mutants had variable *Nodal* patterns: left-sided (2/8), bilateral (3/8) or right-sided (1/8) (Fig. 4A-C). Whereas left-sided patterns were similar to controls, bilateral or right-sided patterns were less propagated along the anterior-posterior axis, failing to expand posterior to the node, and reaching less anterior levels. Some embryos were collected immediately after imaging of the node flow, showing that pattern variability did not reflect detectable flow differences (Fig. 4B2-B4, C2-C4). This rather indicates that the laterality of *Nodal* is variable in the absence of node flow, but still patterned in stripes of lateral plate mesoderm.

**Fig. 4.**
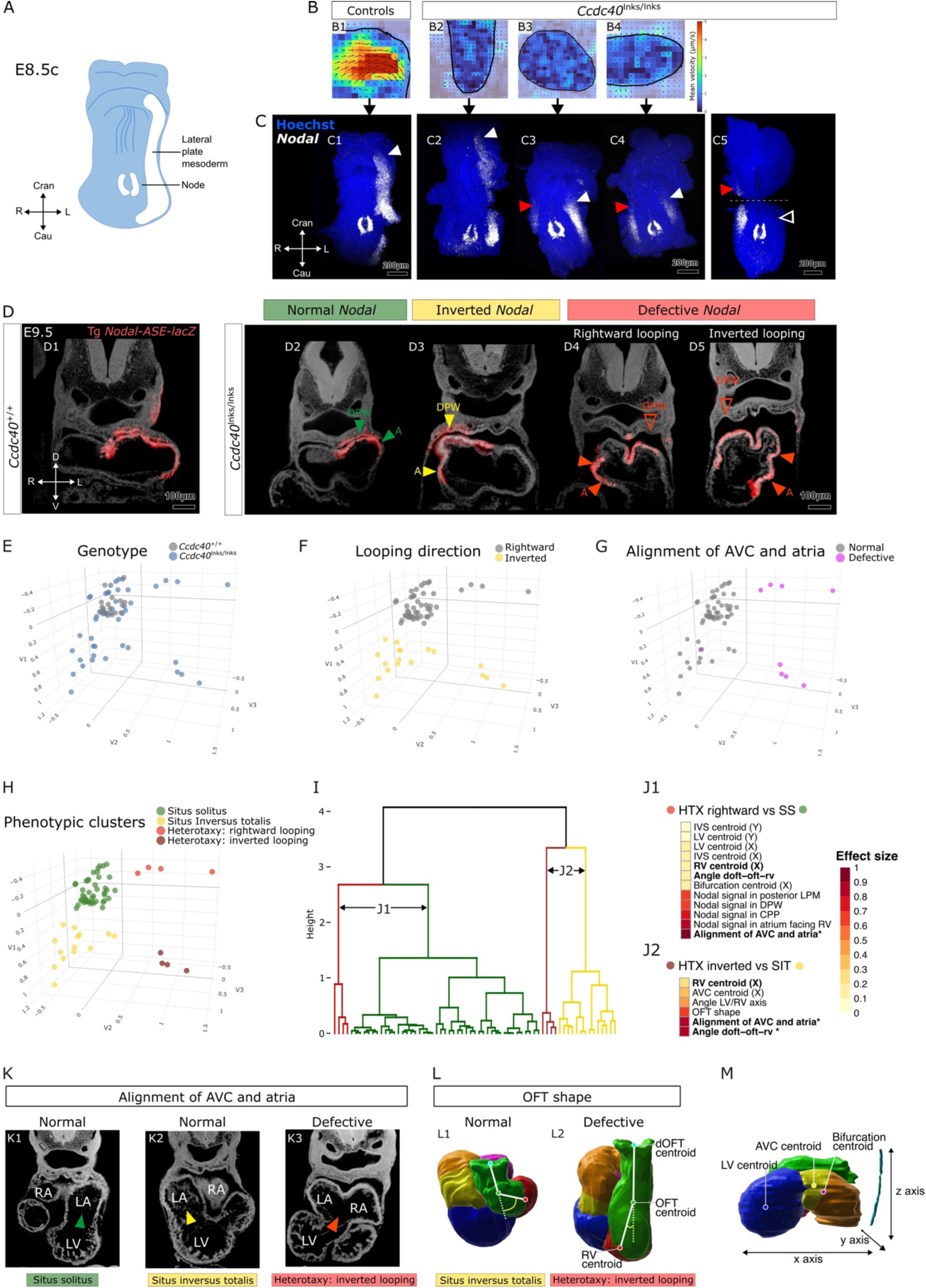
Clustering of *Nodal* pattern in *Ccdc40* mutants. (**A**) Schematic representation of *Nodal* expression (white) at E8.5c. (**B-C**) Correlated analysis of embryos at E8.5c, showing *Nodal* expression by whole-mount in situ hybridisation (white in C), after imaging of node flow (B) as in Fig.1H-I. Genotypes are indicated above. The anterior limit of *Nodal* expression in the left and right lateral plate mesoderm is highlighted with white and red arrowheads, respectively. The embryo in C5 was not assessed for node flow and is shown in separate anterior and posterior views. (**D**) Longitudinal analysis of *Nodal* expression by genetic tracing at E9.5. Transverse sections of *Ccdc40^+/+^; Tg Nodal-ASE-lacZ* (D1) or *Ccdc40^lnks/lnks^; Tg Nodal-ASE-lacZ* (D2-D5) embryos imaged in 3D by HREM are shown. Beta-galactosidase staining is in red and histology in grey. Examples are shown of mutants with a normal (D2, n=15), mirror-image (D3, n=4), or defective (D4-D5, n=3) distribution of cells which have expressed *Nodal*, in the dorsal pericardial wall (DPW) and atria (A). (**E-H**) Unsupervised clustering of a collection of 67 E9.5 embryos, based on 43 geometrical parameters extracted from the 3D segmentation of the looped heart tube, and 5 domains of expression of the *Nodal-ASE-lacZ* transgene. Axes correspond to the PLS regression components. The genotype of embryos is mapped in (E), their heart looping direction in (F), their alignment between the atrioventricular canal and atria in (G) and their cluster identity in (H). (**I**) Corresponding clustering tree, including 17 controls and 28 mutants with situs solitus, 14 mutants with situs inversus totalis, 4 mutants with heterotaxy and rightward looping, 4 mutants with heterotaxy and inverted looping. (**J**) Parameters significantly associated with heterotaxy ordered by effect size, in pairwise comparisons between situs solitus (controls and mutants) and heterotaxy with rightward looping (J1) or between situs inversus totalis and heterotaxy with inverted looping (J2). Common parameters between J1 and J2 are in bold. (**K**) Illustration of the abnormal medial connection of the atrioventricular canal to both atria in a *Ccdc40* mutant in the heterotaxy cluster (K3), compared to a single left atrium connection in situs solitus (K1) and situs inversus totalis (K2). (**L**) Illustration of the abnormal shape of the outflow tract, longer and straighter, in a *Ccdc40* mutant in the heterotaxy cluster (L2), compared to situs solitus (L1). (**M**) Representation of centroids, referred to in (J). A, atrium; AVC, atrioventricular canal; Cran, cranial; Cau, caudal; CPP, cardiopulmonary progenitors; D, dorsal; doft, distal outflow tract; DPW, dorsal pericardial wall; HTX, heterotaxy; IVS, interventricular septum; L, left; LA, left atrium; LPM, lateral plate mesoderm; LV, left ventricle; OFT, outflow tract; R, right; RA, right atrium; RV, right ventricle; SIT, situs inversus totalis; SS, situs solitus; V, ventral. See also Table S3 and Movie S4.

In some embryos, both controls (1/5) and mutants (2/8), we did not detect any *Nodal* expression in the lateral plate mesoderm, whereas *Nodal* was expressed in the node. Since *Nodal* is very transiently expressed (*9*), this may reflect a too early stage of observation. To overcome the sensitive timing of *Nodal* expression and further correlate *Nodal* pattern with morphogenesis, we took advantage of the *Nodal-ASE-lacZ* transgenic line, in which beta-galactosidase is produced under the control of the asymmetric enhancer of *Nodal* (*42*). Because of the perdurance of the *lacZ* mRNA and/or beta-galactosidase protein, the transgenic pattern is still detectable after endogenous *Nodal* expression has been turned off (*9*). We crossed the transgene into *Ccd40* mutants and dissected embryos at E9.5, to correlate the shape of heart looping with *Nodal* pattern. 11/23 *Ccdc40* mutants were indistinguishable from wild-types. They had a rightward looped heart tube with *Nodal* signal in the inferior-left outflow tract, superior-right atrio-ventricular canal, and dorsal left atrium. *Nodal* signal was also seen in heart precursors in the left dorsal pericardial wall and left cardiopulmonary progenitors (Fig. 4D1-D2, Tables S2-S3, Movie S4). *Nodal* signal further expands posterior to the heart in the left lateral plate mesoderm. In 2/23 mutants with an inverted direction of heart looping, *Nodal* signal appeared as a mirror-image (Fig. 4D3, Tables S2-S3). However, the posterior lateral plate mesoderm had bilateral *Nodal* signal. Finally, 9/23 mutants, with either a rightward or inverted looping direction, had more complex patterns of *Nodal* signal, abnormally inferior or absent in the atrioventricular canal, absent in the dorsal pericardial wall, and bilateral in the atria or cardiopulmonary progenitors (Fig. 4D4-D5, Tables S2-S3). Overall, our longitudinal tracking of *Nodal* pattern at E9.5 supports the existence of three categories of *Ccdc40* mutants: normal, complete mirror-image and defective.

We noticed that the defects in geometrical parameters of the looped heart tube and in the laterality of specific domains of *Nodal* signal did not always correlate. We thus decided to take a global multivariate approach and performed unsupervised clustering of embryos, based on their variable parameters (Table S3), resulting in a morphospace with four groups (Fig. 4E-I). One cluster associates together wild-types and *Ccdc40* mutants with rightward heart looping, thus corresponding to the situs solitus phenotype. One cluster contains mutants with inverted heart looping and mainly normal atrioventricular canal, corresponding to situs inversus totalis. The remaining clusters of mutants have consistent abnormal atrioventricular canal, corresponding to heterotaxy, subdivided into either rightward or inverted heart looping direction. To identify the parameters which are most important to define heterotaxy, we performed pairwise comparisons of clusters with the same heart looping direction. We found that *Nodal* signal in the dorsal pericardial wall, cardiopulmonary progenitors, atria and posterior lateral plate mesoderm, as well as the alignment of the atrioventricular canal and atria, the outflow tract shape and ventricle position were the most significant parameters associated with heterotaxy (Fig. 4J-M). These features are reminiscent of heterotaxy defects at birth in the great arteries, complete atrioventricular septal defects and left atrial isomerism (Fig. 2E-F, Table S1).

Taken together, our analysis of heart looping at E9.5 shows the origin of the categories of laterality phenotypes at birth: situs solitus, situs inversus totalis and heterotaxy. In addition to the direction of heart looping, the pattern of mesoderm cells which have expressed *Nodal*, as well as the geometry of the outflow tract, atrioventricular canal and position of ventricles appear as key factors to cluster phenotypes.

### *Nodal* laterality in the lateral plate mesoderm and node crown do not correlate in *Ccdc40* mutants

The lateralisation of *Nodal* in the lateral plate mesoderm was shown to depend on that in the node crown (*8,20*). We thus wondered whether the variable laterality of *Nodal* in the lateral plate mesoderm of *Ccdc40* mutants reflects laterality in the node crown. We quantified *Nodal* asymmetry in the node crown at E8.5c. In controls, *Nodal* is 1.7 ± 0.3 fold more expressed on the left side (Fig. 5A-B), whereas it becomes symmetrical on average in the absence of flow (Fig. 5B). In addition, the slight expression bias of *Nodal* in mutant nodes did not correlate with the expression side in the lateral plate mesoderm (Fig. 5B-D). This indicates that the laterality of *Nodal* patterning in the lateral plate mesoderm is uncoupled from the node in *Ccdc40* mutants, suggesting that it can be regulated by factors others than the node flow or crown asymmetry.

**Fig. 5.**
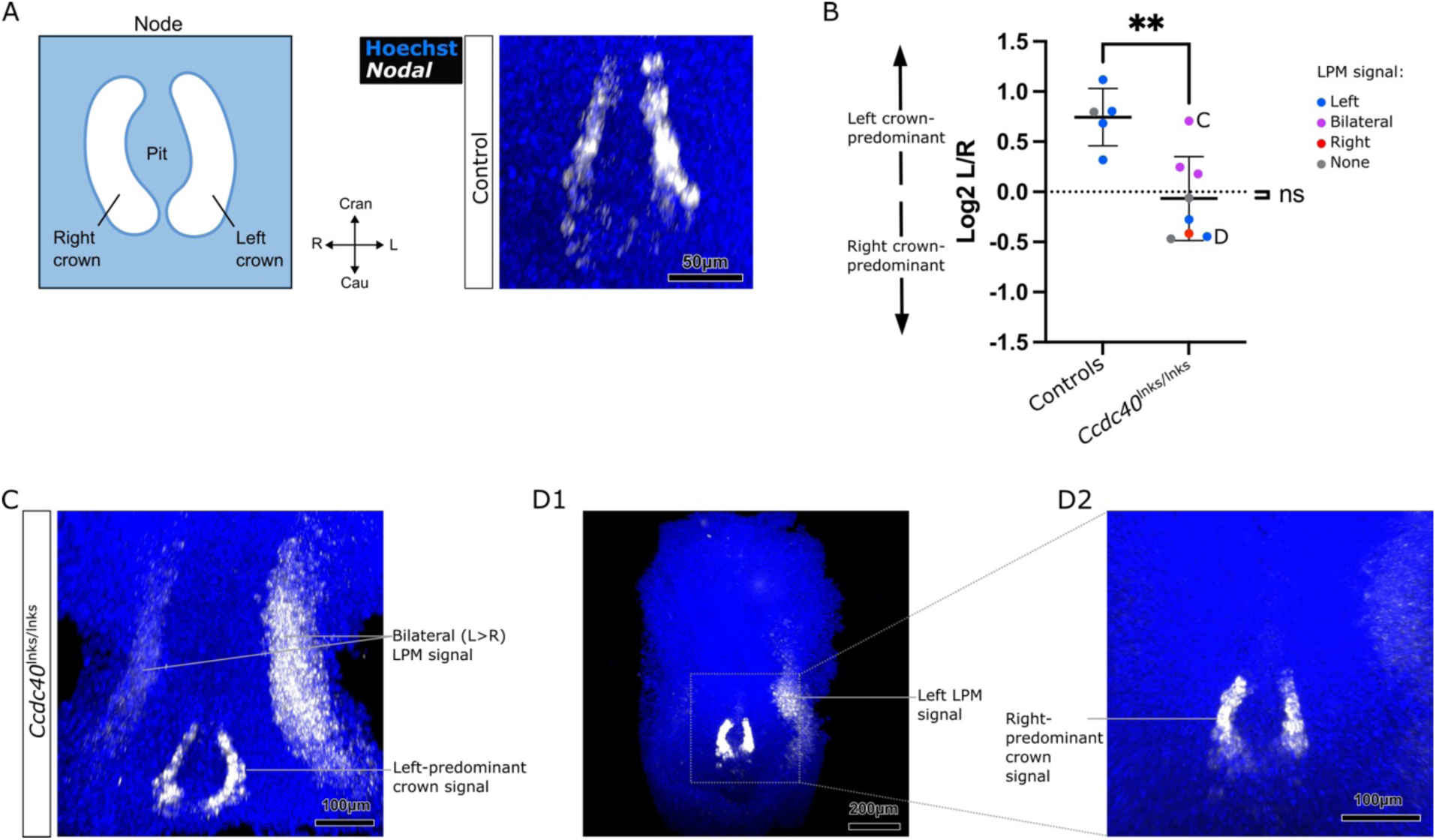
Uncoupling between *Nodal* asymmetry in the node and lateral plate mesoderm in *Ccdc40* mutants. (**A**) Schematic representation and fluorescent in situ hybridization of *Nodal* expression (white) in the node of a control embryo at E8.5c. (**B**) Corresponding quantification of *Nodal* expression in the left versus right node crown, in controls and *Ccdc40* mutants. Datapoints have been coloured according to *Nodal* expression in the lateral plate mesoderm of the same embryo. Means and standard deviations are shown. ** p value=0.006 between controls and mutants (Mann-Whitney test, n=5 control *Ccdc40^+/+^* or *Ccdc40^lnks/+^*embryos, 8 *Ccdc40^lnks/lnks^* mutant embryos). ns, non-significant difference between the mutant mean and a symmetry hypothesis (Log2 ratio =0, One-sample t-test). (**C, D**) Illustration of *Nodal* expression in *Ccdc40* mutants, corresponding to the datapoints indicated in (B). Cran, cranial; Cau, caudal; L, left; LPM, lateral plate mesoderm; R, right. See also Table S6.

### Asymmetry of the WNT pathway and cardiopulmonary progenitors stratify laterality phenotypes

To identify novel asymmetries in *Ccdc40* mutants, we adopted a transcriptomic approach. We micro-dissected the field of heart progenitors at a specific stage (E8.5i) when looping direction can be determined. Paired left and right samples were collected for comparison of gene expression (Fig. 6A). Unsupervised clustering of gene expression asymmetry identifies three groups of embryos (Fig. 6B). As seen in our analysis at E9.5, one cluster associates together wild-types and mutants with rightward heart looping, thus corresponding to the situs solitus phenotype. Another cluster contains mutants with inverted heart looping direction, corresponding to situs inversus totalis. The remaining cluster of mutants corresponds to heterotaxy with rightward heart looping. To extract the main transcriptomic signature of clusters, we performed a differential analysis comparing left and right expression within each laterality cluster. We thus identified a set of 8 genes, which are common and significantly changed both between situs solitus and situs inversus totalis, as well as between situs solitus and heterotaxy (Fig. 6C). One gene, *Ptgfr*, is right-sided in situs solitus, whereas the others, *Pitx1*, *Postn*, *Trabd2b*, *Retreg1*, *Ccn4, Misp3* and *Hspb8* are left-sided.

**Fig. 6.**
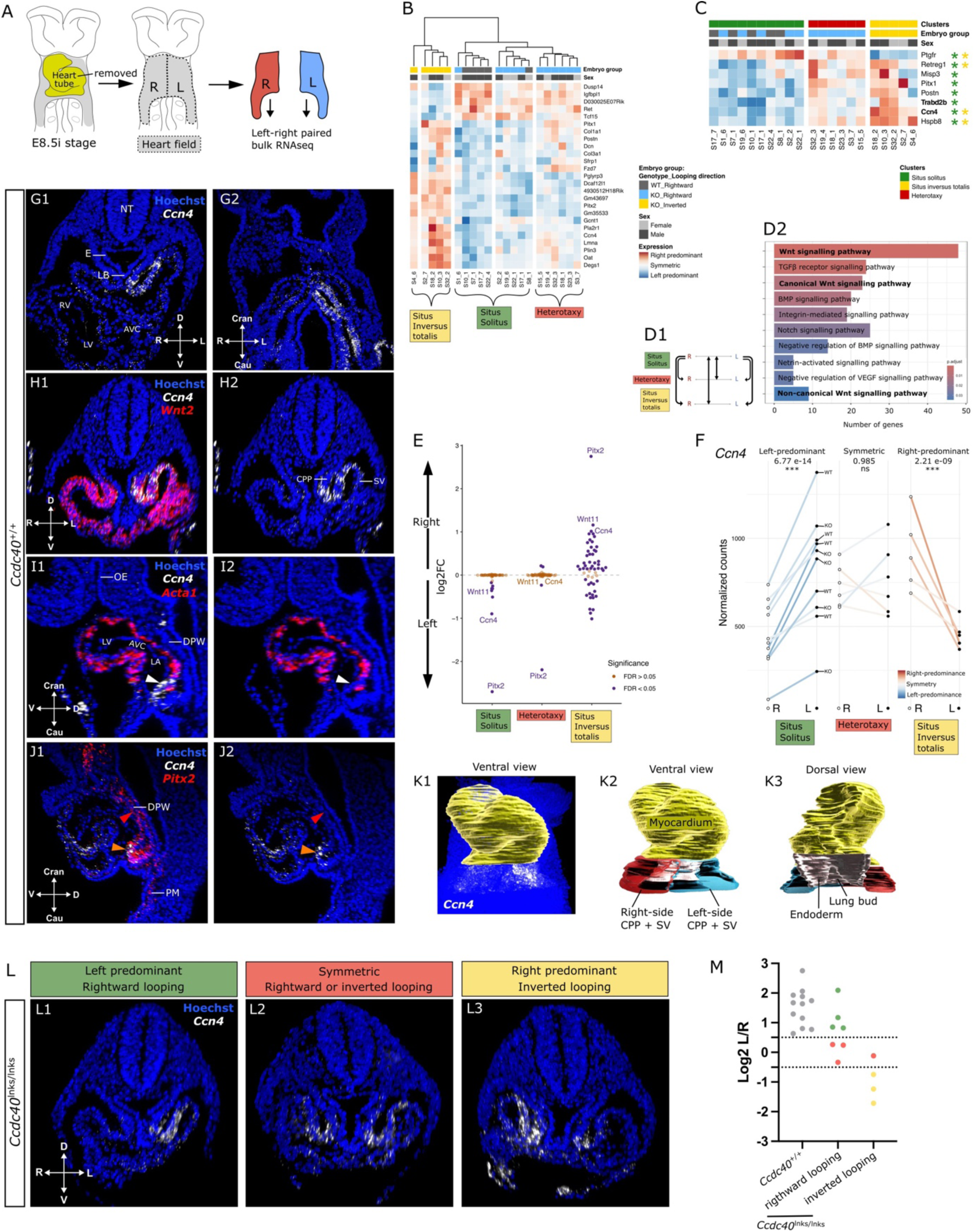
Left-right transcriptomic signature of *Ccdc40* mutant phenotypes at E8.5i. (**A**) Schematic representation of heart field micro-dissection at E8.5i for paired left-right transcriptomics. (**B**) Unsupervised clustering of 21 embryos, based on the 25 most consistent asymmetric genes in wild-type embryos. The genotype, sex and heart looping direction of embryos is colour-coded. Situs solitus includes 5 wild-types and 5 mutants, situs inversus totalis 5 mutants and heterotaxy 6 mutants. (**C**) Transcriptomic signature of clusters. Heatmap of the asymmetric expression of genes which are common and significantly changed in differential analyses between situs solitus and situs inversus totalis clusters and between situs solitus and heterotaxy clusters. Genes associated with Wnt GO terms are in bold. Asterisks indicate significance of asymmetry per cluster (adjusted p-val). (**D**) Schematic representation of the 6 pairwise comparisons between embryo clusters (D1) used to perform a GO analysis of signaling pathway enrichment. Pathways are ordered by decreasing order of significance level. (D2). The most significantly changed pathway is WNT (3 GO terms in bold), representing a total of 56 distinct genes. (**E**) Plot of asymmetric expression of WNT pathway genes (n=56) in the indicated embryo clusters. The WNT pathway is fully left-predominant in situs solitus, largely devoid of asymmetry in heterotaxy and several genes are right-enriched in situs inversus totalis. (**F**) Plot of *Ccn4* expression levels by transcriptomics. The adjusted p-values of right over left expression levels are shown. (**G-J**) Whole-mount in situ hybridisation of *Ccn4* (white) in wild-type embryos at E8.5i, relative to a marker of cardiopulmonary progenitors (*Wnt2*, red in H, n=6), a marker of cardiomyocytes (*Acta1* red in I, n=4), and the left marker *Pitx2* (red in J, n=2). The white arrowhead points to *Ccn4* expression in undifferentiated cardiac cells, the orange arrowhead to overlapping expression of *Ccn4* and *Pitx2* in cardiopulmonary progenitors and the red arrowhead to *Pitx2* only expression in the dorsal pericardial wall. Transverse (G1, H), frontal (G2) and sagittal (I, J) sections are shown. (**K**) 3D segmentation of the *Ccn4*-positive domain relative to the heart tube (yellow) and the foregut endoderm (grey), divided in right-side (red) and left-side (blue). Segmentations are shown overlying the raw in situ hybridisation (K1), in ventral (K1-K2) or dorsal (K3) views. (**L**) Whole-mount *Ccn4* expression (white) in *Ccdc40* mutants at E8.5i, seen in transverse sections, showing three patterns: left-predominance in embryos with a rightward heart looping (L1, n=4), right-predominance in embryos with inverted heart looping (L3, n=3) and symmetric expression (L2, n=4) in the sinus venosus and cardiopulmonary progenitors. (**M**) Corresponding quantification of *Ccn4* expression in the segmented blue/red (left/right) domains shown in (K) in embryos with the indicated genotype and heart looping direction. Abnormal values (red) are outside the 95% distribution interval of control samples based on a Gaussian distribution (dashed lines). AVC, atrioventricular canal; CPP, cardiopulmonary progenitors; Cran, cranial; Cau, caudal; D, dorsal; DPW, dorsal pericardial wall; E, endoderm; FC, fold change; KO, knock-out (*Ccdc40^lnks/lnks^*); L, left; LA, left atrium; LB, lung bud; LV, left ventricle; ns, non-significant; NT, neural tube; OE, oral ectoderm; PM, posterior mesoderm; R, right; RV, Right Ventricle; SV, Sinus Venosus; V, ventral; WT, wild-type. See also Fig. S3-S4 and Tables S4, S6.

Surprisingly, *Pitx2*, a NODAL-target gene was not recovered as a main transcriptomic signature, because it was not differentially sequenced between situs solitus and heterotaxy (Fig. S3A). Although *Nodal* patterning is a key determinant of laterality groups at E9.5, *Pitx2* laterality is difficult to extract by sequencing, because it is entangled with symmetric expression domains (e.g. yolk sac) and several isoforms (Fig. S3B-C). However, transcriptomic changes in the NODAL pathway were detected (Fig. S3D), notably in a right-predominant inhibitor *Ism1*. Overall, in situs inversus totalis, we identified 42 genes with an inverted laterality and a high number gained asymmetry compared to situs solitus (Fig. S3E). One example is *Lefty2*, a NODAL target, which is normally no longer expressed at E8.5i in wild-types. However, it was significantly right-sided in situs inversus totalis (Fig. S3D-E). This suggests that embryos with situs inversus totalis have abnormal kinetics of the NODAL pathway, potentially delayed, as also suggested by the lower antero-posterior propagation of *Nodal* expression in the lateral plate mesoderm at E8.5c (Fig. 4C5). Among genes with gained asymmetry in situs inversus totalis are markers of the atrioventricular canal (*Bmp2*, *Gal*) or the sinoatrial node (*Shox2*). In heterotaxy, 84 genes became symmetric and several gained asymmetry compared to situs solitus (Fig. S3F). Gained asymmetry again probably reflects a delayed developmental programme, given the lower antero-posterior propagation of bilateral *Nodal* expression in the heterotaxy group at E8.5c (Fig. 4C3, C4) and the transient nature of left-right asymmetries (*9,40*). As a user-friendly community resource, we have created a web-interface to interrogate this unique transcriptomic dataset.

We then performed a Gene Ontology (GO) term analysis of genes identified in a broader differential analysis, including also differences in expression levels between embryo clusters (Fig. 6D1). Among the terms reflecting pathways, the WNT pathway emerged as the most significantly affected, representing 56 distinct genes (Fig. 6D2, Table S4). Whereas this set of genes was left-sided in wild-types, it was largely symmetrical in the heterotaxy cluster, and showed either reversed laterality (e.g. *Ccn4*, *Trabd2b*, *Wnt11*, *Pitx2*) or new asymmetry (e.g. *Sfrp2*, *Wnt16*, *Fzd10*) in the situs inversus totalis cluster (Fig. 6E). WNT pathway genes were mainly affected for their laterality (Fig. 6E) rather than their level of expression (Fig. S4E). *Ccn4,* also known as *Wnt inducible signaling pathway protein 1* (*Wisp1*), stood out as the most representative of embryo clusters among genes of the WNT pathway (Fig. 6F): left-sided in situs solitus, right-sided in situs inversus totalis and symmetric in heterotaxy. We further explored the spatial expression pattern of *Ccn4*. We found high expression in a specific domain caudal to the heart tube at E8.5i, covering the sinus venosus and cardiopulmonary progenitors, that face the infolding of the lung bud in the foregut endoderm (Fig. 6G-K). These *Ccn4-*positive cells do not express the myocardial marker *Acta1* (Fig. 6I), but are positive for the cardiopulmonary progenitor marker *Wnt2* (Fig. 6H). *Ccn4* expression is enriched on the left side, overlapping with the left marker *Pitx2* (Fig. 6G, J). *Ccn4* is also expressed in the yolk sac, but symmetrically, and at low levels in the heart tube (Fig. S4A-C). To validate our transcriptomic analysis, we mapped *Ccn4* expression in *Ccdc40* mutants (Fig. 6L-M). We found 4/11 mutants with left-predominant *Ccn4* expression, similar to wild-types. These mutants had a rightward heart looping direction, reflecting the situs solitus cluster. In 3/11 mutants, *Ccn4* expression was reversed, as well as heart looping, reflecting the situs inversus totalis cluster. The remaining 4/11 mutants displayed symmetrical *Ccn4* expression. These mutants had either a rightward or inverted heart looping direction, reflecting the heterotaxy cluster. *Ccn4* thus identifies asymmetry in cardiopulmonary progenitors. Additional markers of cardiopulmonary progenitors can become asymmetric in situs inversus totalis or heterotaxy (Fig. S4D).

Overall, we demonstrate a strong WNT pathway signature of the laterality phenotypes. We identify for the first time an asymmetry in cardiopulmonary progenitors, marked by the WNT pathway gene *Ccn4*, which stratifies phenotypes and is concordant with combined heart and lung defects in heterotaxy and situs inversus totalis.

## Discussion

In a model of motile ciliopathy, we demonstrate that the mouse node is not the only event of symmetry breaking. It is required as a bias for the robustness of left-right patterning, but that asymmetry can be established without any detectable flow. We show the existence of a situs solitus developmental trajectory, with left-sided *Nodal* patterning, normal asymmetric transcriptomics in the heart field, leading to overall normal embryonic morphogenesis and organ laterality at birth, and occasionally mild asymptomatic anomalies. In another developmental trajectory, situs inversus totalis, fully mirror image embryonic morphogenesis and organ laterality at birth arise from right-sided *Nodal* patterning and an inverted asymmetric transcriptomic signature including in the WNT pathway. However, this trajectory shows anomalies in *Nodal* patterning, with temporal delay and posterior bilaterality. Finally, the heterotaxy trajectory manifests in the absence of anterior *Nodal* patterning in the dorsal pericardial wall, bilateral *Nodal* patterning in atria and cardiopulmonary progenitors. It is associated with abnormal symmetry of the WNT pathway, variable direction and defective geometry of heart looping. At birth, this trajectory results in a typical heterotaxy syndrome with left isomerism of the airways and atrial appendages, as well as complex congenital heart defects. Overall, we demonstrate that laterality phenotypes not only correlate with *Nodal*, but also WNT pathway patterning. Our transcriptomics identifies a number of asymmetric genes which characterise these trajectories. We also uncover asymmetry in a cell population relevant to laterality defects, the cardiopulmonary progenitors.

The three categories of laterality phenotypes at birth seen in *Ccdc40* mutants are in line with the phenotype of other mice or patients with motile cilium gene variants (*29,43*). This is completely different from *Nodal* mutants, which have full penetrance of heterotaxy (*9*). Although heart looping direction in *Nodal* mutants is uniformly randomised, their inverted (leftward) looping is not a mirror image and no situs inversus totalis is detected. All *Nodal* mutant fetuses show heterotaxy with right isomerism of the airways and atrial appendages, while heterotaxy occurs with left isomerism in *Ccdc40* mutants. In the clinics, heterotaxy with right or left isomerism tends to be associated with specific congenital heart and spleen defects (*1,44*). Our quantitative analyses of *Ccdc40* and *Nodal* mutants now outline a mechanistic stratification of laterality defects. Human patients with variants in motile cilium genes display laterality defects similar to *Ccdc40* mouse mutants, as well as the respiratory disease primary ciliary dyskinesia and infertility (*26,27,45*). Although the node flow is upstream of NODAL signaling (*6,13*), abrogating node flow leads to a completely different spectrum of defects compared to *Nodal* inactivation.

Using novel tools, we have assessed *Nodal* patterning with higher resolution. *Nodal* is transiently expressed and dynamically propagates spatially (*6,9,46*), the timing of which can be affected in mutants (*28,32,47*). Our genetic tracing of *Nodal* with a transgenic reporter opens the possibility to analyse cumulative NODAL signaling, better catching its overall expression pattern and correlation with embryonic morphogenesis. The sensitive fluorescent in situ hybridization that we have used provides new quantitative insight into asymmetries, compared to previous chromogenic substrates. Our paired transcriptomic analysis shows that situs inversus totalis is not only a phenotypic mirror image of situs solitus, but also a different developmental trajectory, with delayed NODAL signaling. Based on 3D imaging, we have uncovered a cardiac cell population with left-right asymmetry, the cardiopulmonary progenitors. This population had been previously characterised as a pool of common precursors to the heart and lung, marked by WNT signaling components (*48,49*). We now report asymmetry in *Ccn4* and *Nodal* genetic tracing in cardiopulmonary progenitors, which provides a molecular basis to combined heart and lung defects in heterotaxy.

*Nodal* mesoderm patterning in *Ccdc40* mutant embryos is channeled in three types – left-sided, anteriorly right-sided, and abnormal, in agreement with other mutants disrupting the node flow or flow sensing (*6,28,29*). Their frequency is not uniformly randomised in *Ccdc40* mutants, and *Nodal* is spatially not noisy, but still expressed in stripes of lateral plate mesoderm. This raises the question of the origin of the variations. Partial disruption of the node flow can explain phenotypic variability (*30*). However, *Ccdc40* mutants have a fully penetrant undetectable flow, in agreement with the reported immotility of cilia devoid of CCDC40 (*50*). Within the current model of a linear cascade of left-right patterning, a single bilateral *Nodal* pattern would be expected in the absence of node flow, because this prevents the left-sided decay of *Dand5*. *Dand5* is indeed symmetrical in *Ccdc40* mutants (*28*) and other models disrupting flow sensing by *Pkd2* or *Dand5* (*19,51,52*). In the opposite situation, upon *Dand5* inactivation or excessive bilateral decay, *Nodal* displays similar left-sided, right-sided or bilateral variations as in *Ccdc40* mutants (*18,53*). Thus, the node and its readout of *Dand5* asymmetry do not appear to be strictly required to set *Nodal* asymmetry but rather to bias it and make it robustly left-sided. We show uncoupling between *Nodal* asymmetry in the node and in the lateral plate mesoderm in the absence of flow. These observations together support the conclusion that, although *Nodal* asymmetry in the mesoderm is clearly sensitive to the node, it can also be established in the lateral plate mesoderm by a different mechanism.

General principles of symmetry breaking have been proposed, including two main ingredients – a bias which is local and able to orient the asymmetry, and an amplification mechanism which can spontaneously generate asymmetry but with no specific orientation (*54,55*). The bias can rely on molecular chirality, as shown for tubulin, which orients the rotation of cilia and thus determines the direction of flow (*13,56*). Brown and Wolpert (*54*) interpreted situs variations in *Dnah11^iv^* mutant mice as a loss-of-function in biasing, which is in line with our conclusion for *Ccdc40* mutants. In these motile ciliary mutants spontaneous generation of asymmetry seems to rely only on the second ingredient and occurs with a variable orientation. The gold standard of spontaneous amplification mechanisms is the reaction-diffusion process, depending on self-amplifying feedback and long-range inhibition (*57,58*). This mechanism was modelled in the lateral plate mesoderm, based on the activator NODAL and its long-range inhibitor LEFTY (*59*). However, in the absence of *Nodal* in the lateral plate mesoderm, asymmetric morphogenesis is not lost, some heart looping occurs (*9*). This suggests that there are factors other than NODAL participating in symmetry breaking. As a paradigm shift in left-right patterning, we propose that symmetry breaking not only involves the node flow as a bias, but also an amplification mechanism in the lateral plate mesoderm based on NODAL and another pathway (see working model in Fig. S6).

Our paired transcriptomic analyses of the heart field uncover a strong asymmetric signature in the WNT pathway, that stratifies laterality phenotypes. Among the 56 genes significantly changed between laterality groups are *Ccn4*, *Trabd2b* and *Wnt11*. TRABD2B is a metalloprotease that acts as a negative regulator of the Wnt signaling pathway (*60*). The WNT pathway has previously been shown to be asymmetric, with higher *Wnt3* levels in the left node (*33,34*). DAND5 can also antagonise WNT apart from NODAL (*61*). *Wnt3* and *Dand5* indeed act in feedback loops in the node, to reinforce asymmetry (*33*). Whether WNT in the lateral plate mesoderm could also induce NODAL signaling in the absence of flow is an attractive possibility. *Wnt3a* in the posterior node and caudal mesoderm is required for *Nodal* patterning and asymmetric organogenesis (*62*). In the epiblast, WNT signaling is important to induce *Nodal* expression (*63*). In transgenic medaka, ubiquitous activation of WNT1 at 2-4 somites, induces bilateral *Nodal* expression in the lateral plate mesoderm (*64*). Our transcriptomics analyses uncovered other asymmetric markers, such as the NODAL inhibitor *Ism1*, which is broadly expressed in the lateral plate mesoderm (*65,66*). How *Ism1* asymmetry is regulated is unknown. Other candidate pathways in our transcriptomics include prostaglandin signaling or PITX1, but whether they can induce *Nodal* expression remains to be investigated.

Similarly to the mouse, ablation of the left-right organiser or abrogation of flow in other vertebrates is still compatible with normal laterality, with a lower frequency (*67–70*). Additional pathways other than NODAL signaling have been found. In the fish brain, WNT signaling in addition to NODAL is required for the asymmetry of the habenula (*71,72*). In the frog, chick and rabbit, FGF signaling is an important regulator of left-right patterning, upstream of NODAL signaling (*15,73,74*). In the fish and frog, asymmetric cues before the left-right organiser have been observed, including maternal determinants, such as the TGFb factor GDF1 (*75,76*). Such early developmental cues have not yet been reported in mice.

If the node has a biasing role, the long-standing concept of a left-right organizer may be questioned. Blum et al. (*12*) introduced the term of “laterality organising center” to avoid confusion in the use of the term “node” and distinguish the Spemann-Mangold organiser activity from later laterality patterning activity. An organiser is formally defined as a tissue that can both induce and pattern adjacent cells (*77,78*) but the node is now shown to lack the inducing activity in this definition. Essner et al. (*67*) named the orthologous Kupffer’s vesicle in the fish as a transient embryonic “organ of asymmetry”, a term that would be more relevant to the finding that the node is required for asymmetry robustness rather than induction. Overall, our in-depth quantitative analysis of *Ccdc40* mutants provides novel insights into the mechanisms of symmetry breaking in the mouse and the developmental trajectories of laterality defects.

## Materials and Methods

### Animal models

*Ccdc40^lnks/+^* males and females were crossed to generate *Ccdc40^lnks/lnks^* mutant embryos (*25*). The line was kept in a C56Bl6J background. The Tg *Nodal-ASE-lacZ* transgenic line in a mixed genetic background (*42*) was crossed with *Ccdc40^lnks/+^* animals to generate *Ccdc40* mutants carrying the transgene. The *Ccdc40^lnks/+^* mouse line develops faster, so that collection of fetuses the day before birth corresponds to stage E17.5. Heterozygotes *Ccdc40^lnks/+^* had no phenotype and were used together with wild-types as littermate controls in some experiments. In most instances, only wild-types were used as littermate controls. Male and female embryos were collected and used randomly. Embryonic day (E) 0.5 was defined as noon on the day of vaginal plug detection. At E8.5, embryos were staged based on the morphology of the heart (*9,38*) (Fig. S1A). All embryos and fetuses were genotyped by PCR of the yolk sac or tail tip, using primers listed in Table S5.

Mice were housed in individually ventilated cages containing tube shelters and nesting material, at 21°C and 50% humidity, under a 12h light/dark cycle, with food and water ad libitum, in the Laboratory of Animal Experimentation and Transgenesis of the SFR Necker, Imagine Campus, Paris. Animal procedures were approved by the ethical committees of the Université Paris Cité and the French Ministry of Research.

### *Ccdc40* mutant genotyping

For genotyping of *Ccdc40* alleles, forward and reverse primers spanning a region containing the *lnks* C>A point mutation site were designed (Table S5). After PCR amplification, PCR products were subjected to enzymatic restriction by Hpy188I, which differentially finds one and two restriction sites in the mutant and wild-type amplicons, respectively. The expected products are shown in Fig. S5.

### Scanning Electron Microscopy

Embryos were collected at E8.5b-d and fixed overnight in 2.5% glutaraldehyde. After 3 washes in 0.1 M cacodylate buffer pH 7.4, samples were postfixed in 1% osmium tetroxide in 0.1 M cacodylate buffer pH 7.4 for 1 hour. After 3 washes in water, samples were dehydrated in ethanol. Samples were dried in an automated critical point dryer (Leica EM CPD300) and coated with gold/palladium in a Gun ionic evaporator (PEC 681/1). Images were acquired on a JEOL IT700HR scanning electron microscope.

### Beta-galactosidase staining

*Tg Nodal-ASE-lacZ* embryos were collected at E9.5. They were incubated in cold 250mM KCl cardioplegia solution for 1 minute, fixed for 10 minutes in 4% PFA, 5mM EGTA 2mM MgCl2; then permeabilized for 30 minutes in 0.2% NP40 and 0.1% sodium deoxycholate, and finally stained wholemount with X-gal at 37°C in the dark, overnight. After rinsing in PBS, the staining was post-fixed in 4% PFA overnight before 3D imaging by HREM. The transgenic pattern was recorded independently in different domains (Tables S2-S3). 4 regions of the heart tube were considered: the outflow tract (OFT), atrioventricular canal (AVC), atrium facing the right ventricle, and atrium facing the left ventricle. 2 regions correspond to cardiac precursors: dorsal pericardial wall (DPW) and cardiopulmonary progenitors and sinus venosus (CPP). Finally, the lateral plate mesoderm posterior to the heart (posterior LPM) reflects non-cardiac cells.

### Chromogenic RNA in situ hybridization

Embryos were collected at E9.5. After cardioplegia in cold 250mM KCl for 1 minute, embryos were fixed in 4% PFA overnight at 4°C. Fixed embryos were gradually dehydrated in methanol and stored at −20°C until whole mount in situ hybridization. *Wnt11* and *Bmp2* antisense riboprobes (gifts from S. Evans and C. Vesque respectively) were transcribed from plasmids. Signals were detected by alkaline phosphatase (AP)-conjugated anti-DIG antibodies (1/2000), which were revealed with the chromogenic BM purple (magenta) substrate. Samples were post-fixed and imaged in 3D by HREM.

### Fluorescent RNA in situ hybridization

Embryos were collected at E8.5, fixed in 4% PFA overnight at 4°C, gradually dehydrated in methanol and stored at −20°C. Sensitive whole mount RNAscope in situ hybridization was performed using the Mutliplex Fluorescent 426 v2 Assay (Biotechne) and a published protocol (*80*). *Nodal* (436321-C1), *Ccn4/Wisp1* (501921-C1), *Ccdc40* (1054111-C2), *Isl1* (451931-C1), *Pitx2* (412841-C2), *Acta1* (808831-C3) and *Wnt2* (313601-C2) probes were used. Amplification steps were performed using the TSA cyanine5 and cyanine3 amplification kit. Hoechst (1/1000) was used as a nuclear counterstain. Samples were then cleared in R2 CUBIC reagent and embedded in R2 containing agarose. Multi-channel multi-section 16-bit images were acquired with a Z.1 lightsheet microscope and a 20X/1.0 objective.

### Bulk RNA sequencing

Embryos were collected at E8.5 and a ventral brightfield image of each embryo was acquired to record the heart looping stage. Then embryos were micro-dissected, removing the head, back, trunk below the 2^nd^ somite and heart tube (*40*). The remaining tissue, largely consisting of the heart field in the dorsal pericardial wall and down to the second somite, was cut along the midline in two left and right halves, separately collected, and flash-frozen in liquid nitrogen. Samples were then stored at −80°C. Embryos were selected for bulk RNA extraction based on stage (E8.5i), genotype, direction of heart looping and sex (mixed). E8.5i is defined by the tilted position of ventricles, at an angle of about 40° to the antero-posterior axis, with a visible outflow tract and atrioventricular canal (*38*). Given the low penetrance of inverted heart looping (Fig. 3C) and the difficulty to catch a specific transient stage, we had to collect 276 embryos at E8.5 to find 5 samples with inverted heart looping at E8.5i. We selected blindly a twice higher number of mutant samples with rightward heart looping (n=11) and an equivalent group of controls (n=5), for a total cohort of 21 embryos. Total RNA was extracted in TRIzol-Chloroform and purified using the RNeasy Micro Elute Columns kit (QIAGEN) including DNAse treatment. RNA quality and quantity were assessed by capillary electrophoresis using High Sensitivity RNA reagents with the Fragment Analyzer (Agilent Technologies). All RNA Quality Numbers were >9.1. Only paired left/right samples with minimum 10ng of RNA were retained: 5 paired wild-type samples (4 males, 1 female), 11 paired mutant samples with a rightward looping direction (6 males, 5 females), 5 paired mutant samples with an inverted looping direction (3 males, 2 females).

RNAseq libraries were prepared starting from 10 ng of total RNA using the NEBNext Single Cell/Low Input RNA Library Prep Kit for Illumina. The cDNAs produced from the poly-A+ fraction were PCR amplified in 2 steps (9 cycles for the first amplification and 8 for the second amplification). An equimolar pool of the final indexed RNA-Seq libraries was sequenced on an Illumina NovaSeq6000, with paired-end reads of 100 bases and a mean sequencing depth of 80 million reads per sample.

### Live imaging of node flow

Embryos were collected at E8.5b-d in phenol red-free DMEM with 10% Fetal Bovine Serum (culture medium) at 37°C, as established previously (*30,81*). Pre-heated medium and a microscope heating pad were used to control the temperature. Embryos were imaged individually, while the rest of the litter was cultured in the culture medium at 37°C in rolling bottles in a precision incubator (BTC Engineering, Milton, Cambridge, UK). For live-imaging, embryos were mounted on a microscopy glass with a thin layer of silicone glue, flattened with ventral side up and covered by a ∼20uL droplet of culture medium. 0.2 µm green-fluorescent beads were vortexed, diluted 1:10 in culture medium, and 2µL were added on the embryo. Silicone vacuum grease was applied surrounding the embryo, and mounting was closed with a coverslip. Single-channel 16-bit images were acquired using a confocal microscope equipped with Yokogawa CSU-X1 Spinning Disk technology on an inverted Axio Observer.Z1, with an incubating chamber at 37°C, and an ORCA FLASH4 high-speed camera (Hamamatsu). Brightfield images with 10x and 63x/1.46 oil immersion objectives were acquired to record embryo orientation. Then movies on the green fluorescent channel with the 63x/1.46 oil immersion objective were acquired, for 3 focal planes per embryo, at 100fps over 3 seconds. After imaging, 7 embryos were detached by gentle pipetting and fixed in 4% PFA overnight at 4°C, dehydrated in methanol. They were stored at −20°C and *Nodal* expression was evaluated by fluorescent RNA in situ hybridization.

### Micro-Computed Tomography

Fetuses at E17.5 (n=40) or P0 (n=7) were collected and euthanized by decapitation. As described previously (*41*), the body was immerged in HBSS at 37°C 5min to remove blood, then in cardioplegia solution (110mM NaCl, 16mM KCl,16mM MgCl2, 1.5mM Cacl2, 10mM NaHCO3) at 4°C. Fetuses were fixed in 4% PFA 24h at 4°C, washed and stored in PBS azide. Before imaging, the left forelimb was cut as a lateral landmark, and skin around thorax was removed to facilitate penetration lugol as a contrast agent, incubated for 72h. Images were acquired on a Micro-Computed Tomography Quantum FX (Perkin Elmer). Field of exposure was 10mm diameter, and xyz resolution was 20×20×20μm. After imaging, fetuses were recovered and lugol was washed out. The heart was explanted for HREM 3D imaging.

### High Resolution Episcopic Microscopy (HREM)

Whole-embryos (E9.5) or explanted hearts (E17.5-P0) were embedded in methacrylate resin (JB4) containing eosin and acridine orange as contrast agents (*38,41*). The resin block was sectioned with a microtome, and images of its surface were acquired using the optical high-resolution episcopic microscope and an 1X Apo objective, repeatedly. The tissue architecture was imaged with a GFP filter and chromogenic precipitates with an RFP filter. E9.5 embryo datasets include 720-1765 images with of 1-2.1μm resolution for all three (xyz) dimensions. E17.5-P0 heart datasets include 1000-1700 images, with 2.6-3.8μm resolution for the three dimensions. Occasionally, one slice may be lost or overexposed, creating a minor variation in the overall 3D stack. Icy (*93*) and Fiji (ImageJ) (*94*) softwares were used to crop and scale the datasets. 3D reconstructions and analysis were performed using Imaris. Figure panels correspond to optical sections in the most appropriate plane, i.e. not necessarily in the plane of imaging.

### Phenotyping at perinatal stages

Laterality defects in thoracic and abdominal organs were phenotyped based on micro-CT scans, including bronchi, lungs, heart, venous return to the heart, aortic arch, stomach, spleen, liver and colon (*41*). Cardiac anatomy was evaluated in HREM images, based on the segmental approach (*82*) and IPCCC ICD-11 clinical code. Phenotyping was conducted by two independent observers, one developmental biologist and one cardiac pediatrician. The position of the heart apex corresponds to that of the apex of the left ventricle within the thoracic cavity. The anatomical right and left ventricles are defined based on the presence and absence of a septal attachment of the atrioventricular valve, respectively. The position of ventricles reflects the congruent (D-Loop) or not (L-Loop) position of the anatomic right ventricle on the right side of the left ventricle. Malposition of the great arteries was diagnosed in a transverse plane, based on the position of the aortic valve relative to the pulmonary valve. The ventriculo-arterial connection reflects the position of the great arteries, scored as abnormal in the following two cases: double-outlet right ventricle, when more than half of both arterial trunks arise from the anatomic right ventricle; transposition of the great arteries when the aorta arises from the anatomic right ventricle and the pulmonary trunk from the anatomic left ventricle. The anatomic right and left atria are defined based on the presence and absence of pectinate muscles in the posterior wall of the atrial chamber, respectively. The anatomic right and left lungs are defined based on the number of lobes (four on the right, one on the left). The anatomic right and left bronchi are defined based on their subdivision (early on the right, late one on the left). Heterotaxy was defined according to the criteria of Lin et al (*1*).

### Quantification of heart looping at E9.5

HREM images were used to segment the different compartments of the heart tube using Imaris software. Segmentations were done following histological criteria (Video S3), such as cushion boundaries or the interventricular or interatrial sulcus, aided by the expression patterns of *Wnt11* and *Bmp2*, as described previously (*9*). Nine landmarks along the tube were extracted and used for quantifications, of which 3 are two-dimensional polygonal cross-sections of the tube, and 6 are three-dimensional segmented volumes. Polygons were placed at the distal exit of the outflow tract (doft), at the interventricular sulcus (IVS), and at the point of bifurcation into the two atria. The segmented 3D structures are outflow tract (OFT), right ventricle (RV), left ventricle (LV), atrioventricular canal (AVC), left atrium (LA) and right atrium (RA), of which a centroid and volume were extracted. The angles between three consecutive points along the tube axis were taken. Heart shapes were aligned in 3D using an in-house MATLAB code (*80*) (DOI 10.5281/zenodo.8265324) so that the Z-axis corresponds to the notochord and the X-axis to a perpendicular from the neural tube midline to a ventral point. The centroid of the distal outflow tract is set as the center of reference axes. The orientation of the axis between the left and right ventricles, the displacement of the atrioventricular canal (taken here at the bifurcation point) from the notochord and the tube length were measured as previously (*9*). Abnormal values were determined as falling outside the 95% distribution interval of control samples. Deviation in the shape of the outflow tract in mutants was evaluated qualitatively, based on length and curvature. The alignment of the atrioventricular canal and atria was evaluated qualitatively: alignment with the left atrium is the wild-type configuration, partial alignment with the right, in addition to the left atrium, is defective.

### Quantification of fluorescent in situ hybridization signal

3D images were analyzed using Imaris software for the manual segmentation of expression domains. Landmarks for segmentation of the node crown were set by *Nodal*. For cardiopulmonary progenitors and sinus venosus, segmentation was set in the bilateral horns, from the posterior end of atria to the junction with the common cardinal veins. Differentiated myocardium (marked by *Acta1* or histology), proepicardium, and endocardium were not included in the cardiopulmonary progenitor region. After segmentation, the ‘Spots’ tool in Imaris was used, with the ‘Different spot sizes’ feature, to detect fluorescent signal. The volume of the detected spots was extracted and normalized by the total segmented volume and by Hoechst intensity. The resulting values were used for the calculation of the ratio between the left-side and right-side expression.

### Quantification of node cilia phenotype

For quantification of the frequency of ciliary ballooning, the ‘Cell Counter’ plugin in Fiji was used, and a manual marker placed in each cilium, classifying them as the ‘normal’ or ‘ballooned’ type. For delimitation of concentric zones in the node, GIMP (GNU Image Manipulation Program) software was used. The number of ballooned cilia, divided by the total number of cilia, per zone, was calculated for each embryo.

### Particle image velocimetry analysis

Live-imaging of the node produced stacks of 220 to 280 images. Frames were individually processed in Fiji: background illumination was subtracted by applying the ‘Gaussian blur’ filter and subtracting, then salt-and-pepper noise was removed with the ‘Despeckle’ plugin. Image stacks were analysed using the particle-image-velocimetry software (*83*) on interrogation windows with size 48×48 pixels. Flow maps were computed by averaging the flow over the duration of image acquisition. Node areas were selected manually based on the visible distribution of fluorescent beads within a focal plane in the pit.

### Bioinformatics analyses of published single cell RNA sequences

A list of 114 genes associated with motile cilia was assembled based on literature review (*84–86*). Two single-cell transcriptomic datasets were used for the analysis. The expression and annotation data (cluster and meta-cell identifiers) of whole embryos at E6.5–E8.25 (*79*) were downloaded from GEO (GSE169210) according to https://github.com/tanaylab/embflow. The analysis was conducted at the meta-cell level. The expression and annotation data of microdissected heart regions (*39*) were accessed from https://marionilab.cruk.cam.ac.uk/heartAtlas/ and analyzed at the cluster level, focusing on cardiac clusters Me2 to Me7. Each dataset was normalized independently using voom transformation from limma R package (*87*), centered by gene using the mean of background genes defined as genes not in the motile cilia gene list, scaled by gene, and the two datasets were merged. Motile cilia genes were then ranked by total median expression across meta-cells and cells, and k-means clustering was used to identify three distinct clusters. Expression levels were plotted by boxplots, showing the median, 25^th^ (Q1), 75th quantiles (Q3), and whiskers corresponding to Q1 - 1.5(Q3-Q1) and Q3 + 1.5(Q3-Q1). The number of metacells used is indicated in Table S6.

### Bioinformatic analyses of paired bulk RNA sequences

After demultiplexing, reads were trimmed to remove adapter sequences from the first amplification step. Sequence quality was assessed using FastQC and Picard metrics. Duplicates were removed. FASTQ files were mapped to the ENSEMBL Mouse GRCm39 reference genome using the Illumina STAR aligner and counted by featureCounts from the Subread package (v2.0.6). The resulting sequencing depth was 44 million reads in average. Genes with low expression levels were filtered out, using the filterByExpr function of edgeR R package (*88*) with parameters min.count=5, min.total.count=10, large.n=50 and min.prop=0.2. A principal components analysis was conducted on Variance Stabilizing Transformation (VST) transformed data and on the 500 most variant genes to detect potential outliers, no outlier was detected. All differential analysis were carried out using the DESeq2 R package (*89*), with local fit type for dispersion estimation and ashr type (<u>10.32614/CRAN.package.ashr</u>) to shrink the log2 fold changes. To infer embryo clusters, an initial differential analysis was performed on wild-type samples, considering both sexes together and males independently, to identify genes associated with asymmetry in wild type samples. The DESeq2 model was adjusted for the effect of the side (left/right) and embryo identifier to account for pairing. The resulting lists of differentially expressed genes (DEG) were merged and filtered to remove genes not expressed in myocardial lineages (using published single cell transcriptomics (*39*), we kept genes that belong to clusters Me3-Me7 and removed genes which exclusively belong to Me1-Me2, Ec1-Ec2, En1-En2). This list of markers was refined by retaining “congruent” genes, defined as those exhibiting the same left-right trend across all samples. This refinement resulted in 25 genes, which were used to perform hierarchical clustering of all samples of the dataset, based on Euclidean distance and the ward.D2 aggregation criterion. This retrieved 3 laterality clusters.

A second differential analysis was then conducted to assess (i) asymmetry variations within clusters, using a DESeq2 model adjusted for the effect of the side and embryo identifier, and (ii) to evaluate expression changes within each side, using a DESeq2 model adjusted for the cluster and sex. The goodness of fit of these DEseq2 models was assessed by checking the distribution of the raw p-values, ensuring in particular that the variability of embryos within clusters is lower than the variability between clusters. DEG were filtered to exclude genes not expressed in myocardial lineages. For (i), genes were classified as gained asymmetry compared to the situs solitus cluster, when they had a significant right to left expression in a mutant laterality cluster, an absolute log2 fold change higher than 0.5 (for heterotaxy) or 1 (for situs inversus totalis), at least 200 reads in either side, whereas they had a non-significant right to left expression in situs solitus, with an absolute log2 fold change lower than 0.1. Genes were classified as conserved or reversed asymmetry compared to the situs solitus cluster, when they had a significant right to left expression in a mutant laterality cluster, an absolute log2 fold change higher than 0.5, and at least 200 reads in either side. Genes were classified as lost asymmetry compared to the situs solitus cluster, when they had a non-significant right to left expression in a mutant laterality cluster with an absolute log2 fold change lower than 0.1, whereas they had a significant right to left expression in situs solitus, with an absolute log2 fold change higher than 0.5 and at least 200 reads in either side. We selected as signature genes, the genes which were common and changed both between situs solitus and heterotaxy and between situs solitus and situs inversus totalis.

For a global analysis, we compiled (i) and (ii), corresponding to 1842 genes exhibiting diverse variation patterns (Fig. 6D1), filtered based on significance but not fold change or number of reads. The list includes genes that are symmetric in situs solitus but with altered overall expression levels in abnormal laterality clusters; genes that are asymmetric in situs solitus and show changes in overall expression levels in abnormal laterality clusters; and genes exhibiting an increase, decrease, loss or gain of asymmetry in abnormal laterality clusters. Over-representation analysis was performed on these genes using the enricher function from the clusterProfiler R package (*90*). The analysis was applied to several genes set collections, including the full KEGG collection and its subset filtered for “signaling pathways,” the full collection of GO biological process terms and its subset filtered for “signaling pathways,” and additional collections from Biocarta, Reactome, and WikiPathways. Heatmaps were generated using the VST-transformed count differences between the right and left side, with gene scaling to a standard deviation of 1. Hierarchical clustering of genes was conducted with Euclidean distance and the Ward.D2 aggregation criterion. For Fig. 6B, a global clustering tree of all samples was performed. In other heatmaps, the order of laterality clusters was preserved, with sample clustering performed independently within each laterality cluster.

In the analysis of *Pitx2* isoforms, Kallisto (version 0.51.1) was employed to quantify transcript abundances and identify expressed isoforms. The reference genome used for this analysis was Mus musculus mm10 (GRCm39, release 104). Analyses were conducted using the default parameters of Kallisto.

### Extraction of *Ccdc40* sequence in bulk RNA sequences

To identify variants in the specific *Ccdc40* genomic region, we performed variant calling using samtools version 1.18. The command bcftools mpileup was applied on sequence alignment data stored in BAM format. These BAM files were obtained by cleaning fastq files of adapter sequences and low-quality sequences using cutadapt. Only sequences at least 25 nt in length were considered for further analysis (options -O 6 –trim-n –max-n 1). STAR, with options (–outFilterMismatchNoverLmax 0.05 –outSAMunmapped Within –sjdbOverhang 250 and 2-pass mode by-sample), was used for alignment on the reference genome and get BAM files. The reference genome used for this analysis was the Mus musculus mm10 (GRCm39, release 104). The analysis was restricted to chromosome 11, focusing on the genomic position range 119,141,000-119,143,000. Variant calling was then performed using bcftools call -mv, which applies a multiallelic variant detection model. This produced a VCF file containing the identified variants in the specified region for subsequent analysis: no genetic variation was found in any wild-type samples, whereas the A genetic variation was detected in 57 mutant samples (Fig. S5B). Genetic variation from 5 mutant samples could not be retrieved due to the absence of reads mapping to the position of interest but since the samples are paired, we were able to retrieve the genetic variation for each mutant embryo as mpileup detected variant for at least left or right sample.

### Clustering of E9.5 phenotypes

The dataset was divided into two cohorts (Table S3): one consisting of 28 embryos with complete data (*Ccdc40^lnks/lnks^; Tg Nodal-ASE-lacZ* mutants and littermate transgenic controls) and another of 67 embryos, including additional 39 samples with incomplete data, because they were non-transgenic and thus the *Nodal* transgenic signal was missing. In a PLS analysis, we identified two blocks of variables: one comprising 25 geometrical parameters, computed from measures in 3D segmentations of the heart tube, and the other with 5 *Nodal* signal markers (excluding markers which were not variable or the OFT signal which was redundant with the looping direction). Given the mixed nature of the geometrical variables and the dataset structure, we carried out a Multiple Factorial Analysis (MFA) as an appropriate multivariate method. The geometrical data were organized into five blocks of quantitative variables: coordinates (3D coordinates of 8 reference points), volumes (6 cardiac segments), angles (7 between 3 consecutive reference points, and the orientation of the left and right ventricles relative to the notochord), tube length, absolute value of AVC lateral displacement; and nine blocks of qualitative variables, including looping direction, OFT shape, alignment of AVC and atria, and Nodal signal in different domains (DPW signal, AVC signal, signal in the atrium facing the right ventricle, posterior LPM signal and CPP signal). To account for dependencies between the X, Y, and Z coordinates in the “coordinate” block, the data were summarized using six factors from the DISTATIS method, using the DistatisR R package (<u>10.32614/CRAN.package.DistatisR</u>). MFA for the 67 samples was performed on the geometrical variables, while a Multiple Correspondence Analysis (MCA) for the 28 samples was applied to the *Nodal* signal variables due to their qualitative nature. Both analyses were performed using the FactoMineR R package (*91*). A PLS regression was then conducted to model the relationships between the geometrical and *Nodal* signal markers, enabling us to infer the MCA coordinates for the incomplete cohort of 39 samples. For both the MFA and MCA, we retained enough components to capture 80% of the variability, resulting in 7 and 5 components, respectively. This methodology led to a 3D projection of all 67 samples, with contributions from all variables, including those with missing data, by leveraging the relationships between the geometrical and *Nodal* signal markers. This projection, representing the relations between the two sets of markers, is referred to as the “PLS-space.” PLS regression was performed using RGCAA R package (<u>10.32614/CRAN.package.RGCCA</u>). Hierarchical clustering was applied to the first two PLS components. Marker associations for each cluster were identified through pairwise statistical testing between clusters (e.g., heterotaxy with rightward looping versus situs solitus, heterotaxy with inverted looping versus situs inversus totalis) for all variables. For quantitative variables, the Mann-Whitney-Wilcoxon test was used, while qualitative variables were assessed with Fisher’s exact test. Effect sizes were also calculated: Eta-squared for quantitative variables and Cramer’s V for qualitative variables, both measures implemented in the effectsize R package (*92*).

### Statistical Analyses

Statistical tests and p-values are described in figure legends and Table S6. P-values less than 0.05 were considered statistically significant. Group allocation was based on PCR genotyping and heart looping direction. All sample numbers (n) indicated in the text refer to biological replicates, i.e. different embryos/fetuses or different cells. Investigators were blinded to allocation during imaging and phenotypic analysis, but not during quantifications. To compare whether two groups had equivalent means, a t-test was used when a normal distribution could be assumed, and a Mann-Whitney test otherwise. To test whether the distribution of classes followed an expected distribution (e.g. Mendelian distribution, or random distribution), a chi-square test was used.

## Acknowledgments

We thank L. Guillemot, V. Benhamo, Z. Mercier, J. Terret, N. Agueeff, M. Meunier for technical assistance; F. Corson for the Particle Image Velocimetry algorithm and insightful discussion; T. Holm Bønnelykke, P. Campagne, A. Desgrange, H. Hamada, L. Houyel, J-F. Le Garrec, G. Letort, D. Norris, K. Shinohara, H. Varet, I. Zohn for expert advice; K. Anderson and K. Hadjantonakis for providing *Ccdc40^lnks^*mice; D. Conrozet and the histology platform of the SFR Necker; C. Bole-Feysot of the Genomics platform; N. Cagnard and C. Masson of the Bioinformatics platform; H. Augis Chu and the Cell imaging platform; the LEAT animal facility, the Ultrastructural BioImaging facility of the Institut Pasteur.

## Author contributions

Conceptualization: AO-S, SMM

Methodology: AO-S, EP, SMM

Formal Analysis: AO-S, AC, EP

Investigation: AO-S, ZO, TB, CR

Data Curation: JL

Writing – Original Draft: SMM

Writing – Review & Editing: all authors

Visualization: AO-S, EP, AC

Supervision: AO-S, PB, SMM

Funding Acquisition: AO-S, PB, SMM

